# In the face of competition, imperfect knowledge can lead to sustainable exploitation of a shared resource

**DOI:** 10.1101/2025.01.16.633107

**Authors:** Geert Aarts, John R. Fieberg, Andries Richter, Jan Jaap Poos, Niels T. Hintzen, Allert I. Bijleveld, Geerten M. Hengeveld, Tobias van Kooten, Anieke van Leeuwen, Jason Matthiopoulos, Frank van Langevelde, Louise Riotte-Lambert

## Abstract

Overexploitation of a shared resource is a frequent phenomenon among groups of selfish consumers, known as the ‘tragedy of the commons’. To avoid “self-damage through self-interest”, humans may either defend exclusive access to resources, or collectively adopt advanced cooperative behaviour and carefully designed spatial harvesting strategies to exploit a shared resource sustainably. However, why do many non-human species that are neither social nor territorial also appear to avoid overexploiting resources? This study shows that, even under highly competitive scenarios, cooperation is not necessary to sustainably exploit shared resources if individuals have imperfect knowledge about the distribution of those resources. Poorly informed individuals are more likely to inadvertently exploit resources in peripheral habitats, which may allow for higher resource renewal in the nearby and low-cost habitats, enhancing the sustainability of resource use. As a result, those consumers can achieve a long-term yield that approaches that of cooperative consumers and is higher than omniscient consumers. Ironically, while there might be an evolutionary drive for individuals to make well-informed decisions, consumers of shared resources may be collectively more prosperous when they make decisions based on less information. These findings might explain why apparently sub-optimal consumer behaviour appears so prevalent in nature.

## INTRODUCTION

Many renewable resources are considered common pool resources, meaning they are non-excludable and are subject to competition among consumers. As a result, exploitation by one individual reduces resource availability for others [1]. While it might be beneficial for an individual to delay resource exploitation to allow resources to regrow and achieve higher harvest rates in the future, there is a risk that other individuals will exploit those resources first. This is known as ‘competition by pre-emption’, a mechanism leading to the ‘tragedy of the commons’ [1,2] whereby the exploitation of shared resources by selfish consumers pursuing short-term gains leads to extreme resource depletion. Humans, as a species, have often been directly or indirectly responsible for overexploitation of renewable resources. This has resulted in declines or extinctions of taxa in terrestrial and marine ecosystems worldwide [3,4]; examples include the decline of cetaceans during the whaling era [5], the collapse of cod populations [6], or the extinction of several large vertebrates, like the dodo [7]. The ongoing logging of old-growth forests is an example of overexploitation of plant communities [8]. There are also examples of human-induced overexploitation caused by other species. This can happen when keystone predators are eliminated, resulting in ecological cascades [9–11], or when exotic species are introduced without having native predators [12]. However, it appears that overexploitation is considerably less prevalent in other species than humans [13].

To prevent overexploitation and optimise resource harvest, consumers could carefully consider resource renewal. Resources are commonly assumed to grow in a sigmoidal fashion (the logistic growth model), in which case maximum resource productivity, and hence harvest, can be attained at intermediate resource densities. This idea is also central to the concept of Maximum Sustainable Yield (MSY - [14]), extensively used in fisheries management. Such a strategy, however, falls short when the pay-off for resource harvesting varies over space or time [15,16]. For example, some resources might be more attractive to harvest because they are placed in safer locations (where there is less risk of predation or environmental exposure) or because they are easier to acquire (e.g., less concealed, or more accessible). This is especially the case for central-place foragers who are constrained by their home-base, and who may view nearby resources as being more attractive, leading to local over-exploitation (i.e., exceeding MSY). Indeed, very low resource availability close to the home-base has been observed for many central-place foragers, which is known as Ashmole’s halo [17–20]. To account for these spatial effects, Ling and Milner-Gulland [21] extended the MSY-framework to consider travel costs. They illustrated that social harvesters could attain a higher (sustainable) yield per unit cost by collectively adjusting resource exploitation. This was achieved by considering the shape of the resource renewal function, as well as exploitation and travel costs.

Although social and sustainable exploitation regimes are theoretically appealing, in practice, they are challenging to implement [22–24]. First, they require the harvester to know the underlying exploitation costs, resource density, and the shape of the resource-renewal function. This resource-renewal function is particularly difficult to quantify by the harvester since it requires repeated visits and cues about what other competitors are doing. Second, individuals in competition with others would collectively have to postpone exploitation in the more profitable resource patches nearby and allow resources to replenish, and in the meantime, be prepared to make more costly trips to exploit resources elsewhere. Even when such a strategy (i.e., a delayed reward) would increase average long-term harvest rates for the population, it breaks down in the presence of defectors who would reap the benefits of such local competition release [25,26]. Thus, to avoid the ‘tragedy of the commons,’ individuals would seemingly need to be forced to coordinate their actions and cooperate [27,28], which would require advanced cooperation behaviour. Thus, the emergence of a sustainable exploitation strategy seems to place a large burden on individual cognition and short-term fitness. Instead, could there be a mechanism that enables individuals to act as self-centred consumers, aiming to maximise short-term gains, yet still achieve sustainable exploitation of resources?

One possibility is that long-term optimal exploitation comes about not by collective choice and perfect cooperation, but, rather, by imperfect efficiency in an uncertain and variable world. Previous simulation studies have indeed demonstrated that imperfect information may lead to higher long-term harvest rates [29]. Here, we deepen this insight by examining how consumers’ imperfect perception of resource distribution affects long-term harvest rates, and how this may lead to the emergence of a more sustainable exploitation regime. To do this, we first compare two strategies in the absence of uncertainty: I) selfish individuals that aim to maximise short-term harvest rate, and II) social individuals that can and do consider resource growth and future costs of exploitation, and are willing to incur higher immediate costs for greater future compensation [21]. These strategies, and their corresponding equilibrium resource and consumer population abundances, will serve as reference points of the extremes of a spectrum of possible strategies. We then proceed to examine individuals who prioritise their own interests and maximise short-term harvest rates, but who have imperfect knowledge about the availability of resources. We investigate whether they are less likely to overexploit resources and trigger their collapse, as well as whether they are capable of attaining long-term exploitation rates that are comparable to those of cooperative individuals.

## METHODS

### Resource growth

For living resources, a typical renewal function (*q(x*)) is the logistic growth function:

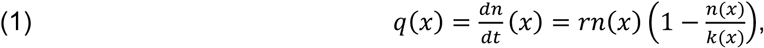

where *n*(x) is the resource density (units/m^2^) at location *x*, *r* is the intrinsic population growth rate of the resource, and *k(x)* is the local carrying capacity density at location *x*. For the moment, we assume that both *r* and *k* are constant throughout space, and we assume no movement in the resource population.

### Resource harvesting strategies

We examine two extreme harvesting strategies: strategy I, which aims to maximise short-term trip-level yield per unit cost; and strategy II, which aims to maximise (long-term) sustainable yield per unit cost. We investigate the effect of each strategy on the equilibrium landscape distribution of resource density and productivity and assess the landscape’s capacity to sustain a certain level of long-term yield. We focus on central-place foragers (CPFs) because they have a simple spatially-varying cost function, namely the travel cost between the home-base and harvest location, represented as a function of distance [21]. Furthermore, we assume that foraging individuals have a set loading capacity (for example, a fixed stomach size), and during each trip only visit a single location in the landscape to collect resources. For these central-place foragers, the total cost (in time) to collect the resources defined as a function of the distance *d* from the home-base is:

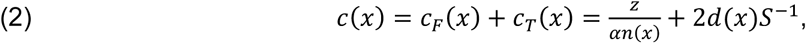

where *c_F_(x)* and *c_T_(x)* are the harvest and travel costs per trip for harvesting at location *x*, respectively. *z* is the (fixed) amount of resources harvested, *α* is the harvest rate parameter (i.e. area harvested per unit of time), *d(x)* is the distance between the home-base and location *x* and *S* is the travel speed.

### Resource exploitation strategy I: Maximizing short-term trip yield per unit cost

Foragers that wish to satisfy their immediate resource needs at the lowest cost should select the patch with the highest resource yield per unit cost for that trip. This corresponds to selecting the patch where the total costs *c(x)* per unit yield (eq. 2) is lowest. Figures 1 and 2 illustrate the decision involved for a discrete patch scenario: An angler must choose between three equal-sized ponds (A, B and C) that differ in travel distance and initial resource density, and thus in travel and foraging times, respectively (Fig. 1). If the angler behaves as a trip harvest-rate maximiser and the costs are time, the angler should choose pond A, since walking to and harvesting from pond A will require the least amount of time per unit fish caught (see Fig. 2a). However, this single decision at the trip level will also have future repercussions. By harvesting, resource density decreases, and consequently, so will resource growth (Fig. 2c). Since the resource density in pond A is below the maximum sustainable yield (MSY), any exploitation will lead to a decrease in resource growth (Fig. 2c).

**Figure 1.**
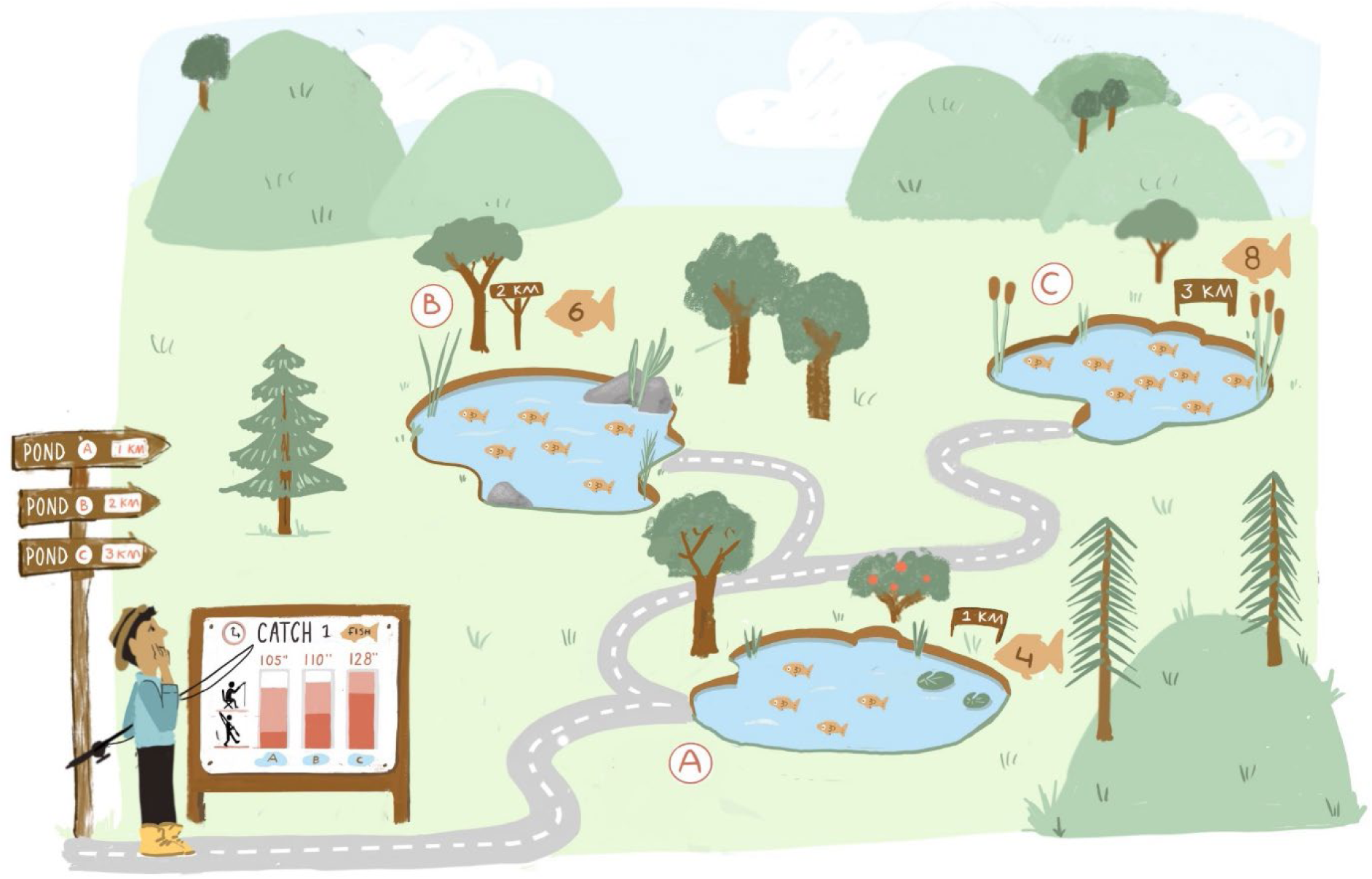
What harvest strategy should be used to catch one fish? An angler has three ponds to choose from (A, B, and C). These ponds are spread out over different distances (1, 2 and 3 km, respectively), and require varying walking times (shown in red on the information sign, in minutes). The ponds also differ in initial fish density, with the highest fish density in pond C (800 fish; each fish symbol represents 100 individuals) and the lowest in pond A (400 fish). Consequently, the fishing times (displayed in pink on the information sign) also differ between ponds. If the angler visits this area only once and wants to maximise the yield (one fish) per unit time, they should choose pond A, as it will take the least amount of time (i.e., 84 minutes in total). However, removing a fish will influence future fish biomass growth and may therefore not be the best decision in the long run.

**Figure 2.**
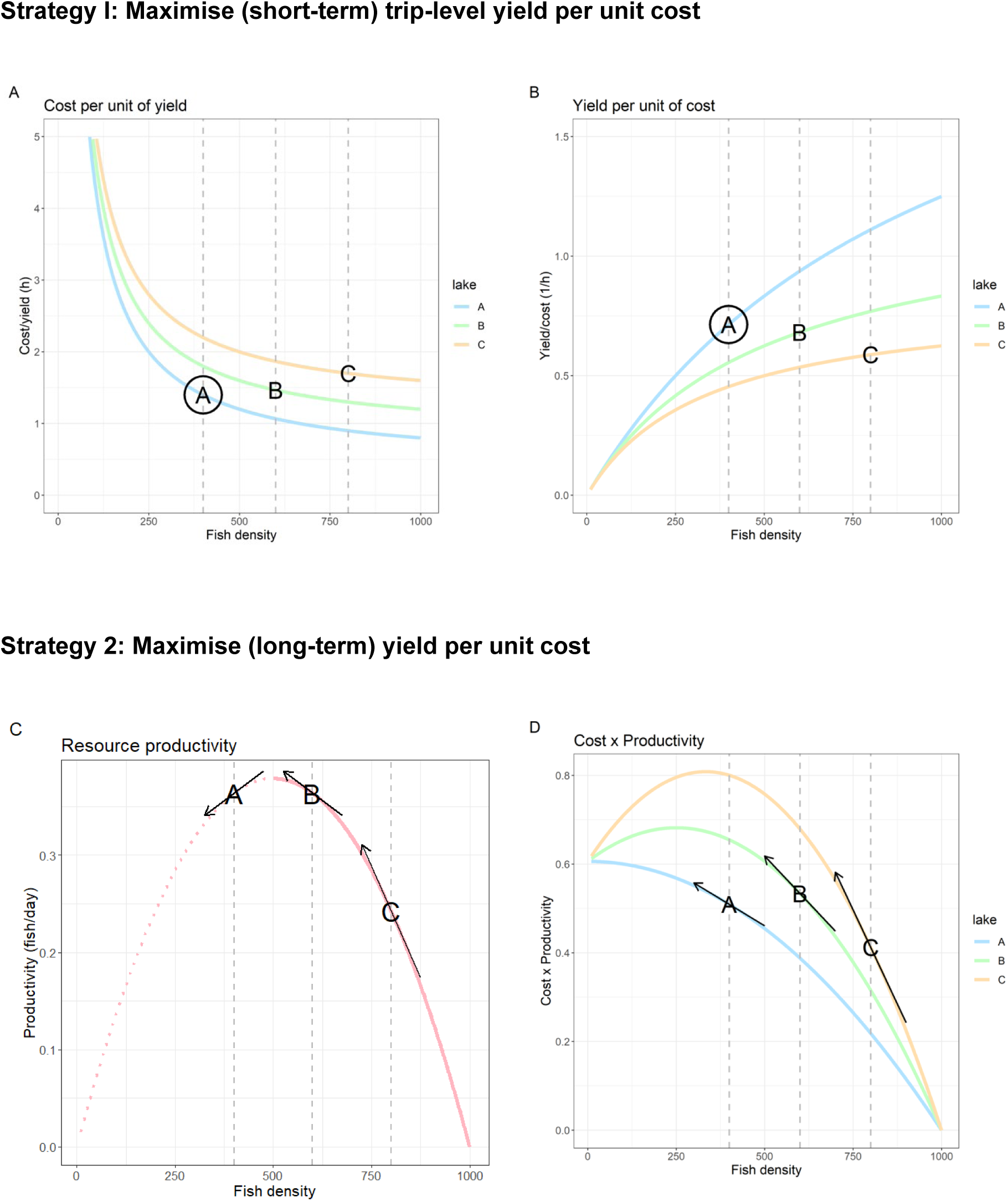
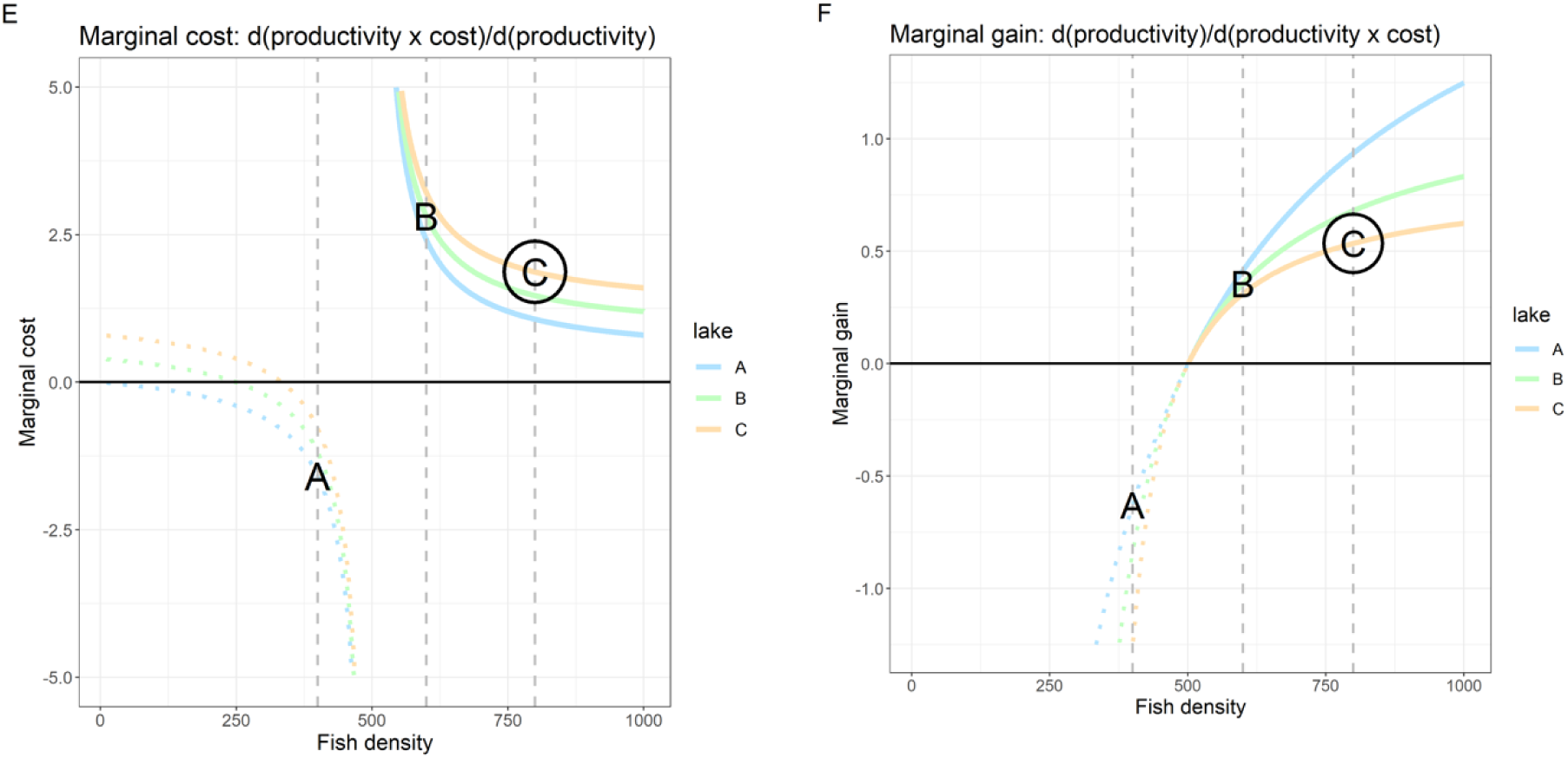
The optimal choice of the three ponds (A, B, or C, see also Fig. 1) depends on whether the (immediate) short-term or long-term yield is maximised. When maximizing the short-term yield per unit cost, the angler should select the pond where the total cost (travel + foraging cost) for harvesting one fish is lowest. Figure 2a shows these costs as function of fish density. At the initial fish densities (A = 400, B = 600 and C = 800), the total cost of harvesting one fish is lowest for pond A (a). Similarly, the yield per unit cost is highest for pond A (b). Hence, for this specific example, a trip-harvest-rate maximiser should select pond A (denoted by black circle). Removing a fish from any of the ponds, however, will have future repercussions. It lowers fish density, and therefore also affects (future) fish renewal. This renewal as a function of fish density (pink line in c) follows a bell-shaped curve, assuming logistic growth with carrying capacity *k* = 1000. The MSY is 500 fish. The arrows indicate the derivative, i.e., the change in renewal after removal of one fish from each pond. After removing a fish from pond B or C, fish renewal increases, while for pond A, fish renewal declines. From the consumer’s perspective, the value of the resource production (shown in c) differs among the three ponds. For example, resource production in pond A, being the closest, is much more valuable since it requires lower travel costs (d). These harvesting costs as function of fish density are shown in d. To obtain the highest long-term yield at the lowest costs, the angler should try to push the system in the direction whereby harvesting leads to the largest increase in productivity and the smallest change in costs to harvest the produced resources (indicated by the arrows in d). The marginal cost is the ratio of the change in costs to harvest the additional resource productivity relative to the change in productivity induced by harvesting at that location (e). The inverse of the marginal cost is the marginal gain (f). A sustainable harvest-rate maximiser will have the highest long-term harvest per unit cost, when selecting the lake with the highest marginal gain. Although the increase in cost to harvest the increase in productivity for lake C is highest, the change in productivity of the system is even higher, and thus, harvesting one fish from lake C results in the largest marginal gain (denoted by black circle).

### Resource exploitation strategy II. Maximizing long-term sustainable harvest per unit cost

When harvesters attempt to maximise the long-term yield per unit cost, they must first carefully consider how harvesting in each pond affects resource growth, since this growth ultimately determines future yield. In the simplified angler example, in ponds B and C, where resource density remains above MSY, harvesting increases resource growth, whereas harvesting in pond A reduces future resource growth. However, consumers must also consider future costs of harvesting the resources produced. These costs are dependent on the location of the pond (which determines travel time) and fish density (which determines the time required to harvest the resource). For example, the cost of harvesting resources in pond C is relatively high due to the extensive travel costs involved. Therefore, any increase in production of resources in pond C is considerably less valuable to the consumers. When consumers attempt to maximise (long-term) sustainable harvest per unit cost they should consider both changes in harvest cost and resource growth, i.e., they should harvest in the location with the largest increase in resource growth (*q(x)*, Fig. 2c) relative to the changes in the cost to harvest these additional resources (*c(x)q(x))*, Fig. 2d). The ratio of these two derivatives is referred to as the marginal gain *F* (Fig. 2f):

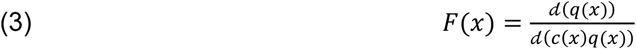

The inverse of the marginal gain is known as the marginal cost [21]. In terms of our pond illustration, while it is most economical to collect resources from the pond A at the trip level (Fig. 2a), it is more beneficial in the long run to first harvest resources from pond C (Fig. 2f).

### Individual-based simulations

Following the short-term exploitation strategy (i.e. strategy I) leads to a lower long-term yield compared to the long-term sustainable exploitation strategy (i.e., strategy II), but how will the relative performance of the two strategies change when information about resource density at the individual level is uncertain? To examine how information uncertainty affects long-term harvest rates, we used individual-based simulations and included stochasticity in the encountered and memorized resource density. We contrasted these results with similar simulations in which individuals possessed perfect knowledge of the density and productivity of the resources (fish, in this instance).

### Simulation single individual

We used the same equations and parameters as outlined in the preceding sections and in Fig. 2. Commencing with a single individual, at the simulation’s onset, this individual first visited each pond once. After completing the walk and reaching the first pond, the angler started fishing until *η* fish (with a total biomass of z) were caught. The ‘waiting’ time required to harvest cryptic resources like fish, is subject to stochasticity and can be conveniently modelled using a Gamma distribution with shape parameter *η* representing the number of resource items to harvest, and the rate parameter μ equal to the product of the catch rate parameter (α, see Appendix A, table A1) and the resource density (*n(x)*) in each pond. After harvesting *η* fish, the angler memorized the time spent fishing, and subsequently returned to their home-base. Overnight, fish were replenished in accordance with the logistic resource renewal function (eq. 1). These fishing trips were repeated for the other two ponds on the following two days. On the fourth and subsequent days, the angler relied on the memorized trip durations of the most recent visits to each pond and selected the pond with the highest anticipated trip yield per unit cost (strategy I, eq. 2). Again, after fishing was completed, the angler memorized the duration of this most recent trip (and erased the memory of the preceding visit) for future decision making.

Due to natural stochasticity, the angler may occasionally take a very long time to catch a fish, which would lead to subsequent avoidance of a specific pond, even if the pond is not inherently resource-poor. To prevent such persistent avoidance, occasional explorations (occurring once every twenty trips) were carried out. In these explorations, a pond was randomly sampled from the ponds with the selection probability inversely proportional to the anticipated total trip duration (including both travelling and harvesting) for each pond.

### Varying information uncertainty by adjusting the level of stochasticity

Information uncertainty arises from the stochasticity in the encounter rate and the fact that individuals are allowed to recall only the most recent visit. Using the Gamma distribution, it is possible to change the variability in harvest duration while keeping the mean duration constant.

One way to accomplish this is to adjust the shape and rate parameters in proportion to each other (Appendix A, Fig. A2). For example, when individuals harvest *η* = 1 resource items at a given resource density, the average harvest duration is *c_f_* and the variability in harvest duration is σ = 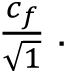 In contrast, when individuals acquire *η* = 1000 smaller resource items when the harvest rate is also 1000 times higher, the average harvest duration *c_f_* to acquire these 1000 resource items remains the same, but the variability in encounter rate is much smaller, namely 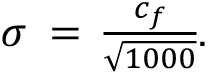 To include different levels of uncertainty, we varied *η* from 1 to 40 000 and adjusted the rate parameter to keep the mean harvest duration constant.

### Simulation multiple individuals

Next, we expanded the simulation to include multiple (m = 50) individuals participating in fishing trips. To achieve the same overall removal as the single individual scenario, each of the *m* individuals harvested only *1/m* resources during each trip. In total, we ran three simulation scenarios. In the first “omniscient” scenario, all individuals followed the long-term maximisation strategy and had perfect (external) information on resource density. In the second “omniscient” scenario individuals also had perfect information but pursued a short-term maximisation strategy. In the third “uncertainty” scenario, individuals followed a short-term maximisation strategy, but they only remembered their most recent visit. Individuals experienced variations in the time required to catch fish (represented using a Gamma distribution with shape parameter *η* = 1) in all three scenarios, but this stochasticity created information uncertainty in only the third scenario. Each simulation was run for 10,000/*m* iterations, during which resource density and yield per unit cost were monitored.

We also examined the impact of a shift in strategy by some individuals on the yield per unit cost. Each simulation scenario was run twice, and after 10 000/*m* iterations, in one simulation run, 50% of the individuals were replaced by individuals using one of the two alternative strategies. For example, for the simulation scenario where individuals followed strategy I and faced uncertainty, 50% of them were replaced by omniscient individuals who adhered to strategy I.

## RESULTS

### Selection of harvest location at the onset of the simulation

An omniscient harvester aiming for short-term gains should choose the pond with the lowest cost per unit of yield (i.e., the highest yield per unit cost). In the illustrated example, at the onset of the simulation, they should therefore choose to harvest in pond A. Although the time spent fishing at A is somewhat longer, this is outweighed by a much shorter travel duration (Fig. 2A, 2B). In the long-term, harvesting at A is a less profitable decision, because the initial resource density at A (400) is significantly lower than the MSY (500), and harvesting will decrease future resource renewal. In contrast, harvesting at B, and particularly at the most distant pond C results in an increase in productivity since the initial resource density is well above MSY (Fig. 2C). Overall, pond C has the highest positive change in productivity (Fig. 2C) relative to the change in cost to harvest this productivity (Fig. 2D), and thus, the largest marginal gain (Fig. 2F). As a result, harvesting at C is the optimal choice for the long-term harvest rate maximiser who takes future productivity into consideration (Fig. 2E, 2F).

### Emerging consequences of the harvesting strategies

Depending on the harvesting strategy, the angler exerts a certain amount of harvest pressure that varies between locations. This leads to differences in resource density between locations, and since resource density determines productivity (eq. 1), resource renewal will also vary among the locations. Eventually, each location converges to an equilibrium resource density where harvest and renewal are in balance. The precise values of these equilibria depend on various parameters – like the resource renewal rate (*r* in eq. 1), fishing pressure (here one fish per day), and fishing efficiency (*α* in eq. 2) – and on the harvesting strategy. If the angler attempts to maximise the short-term yield per unit cost, it will select the pond which requires the least amount of time to catch a fish. If this process is repeated, eventually, in line with the ideal free distribution, the anglers will distribute harvest effort in such a way that the total amount of time required to catch a fish becomes equal across all ponds; 2.0 hours in this example (Fig. 2). Because the travel duration varies between ponds, higher travel duration should be compensated for by lower harvesting duration, and thus, higher resource density. Eventually, the resource density in the closest ponds A and B will be below MSY (i.e., overexploited), while C remains close to the carrying capacity (Fig. 3), where resource renewal is also low.

**Figure 3.**
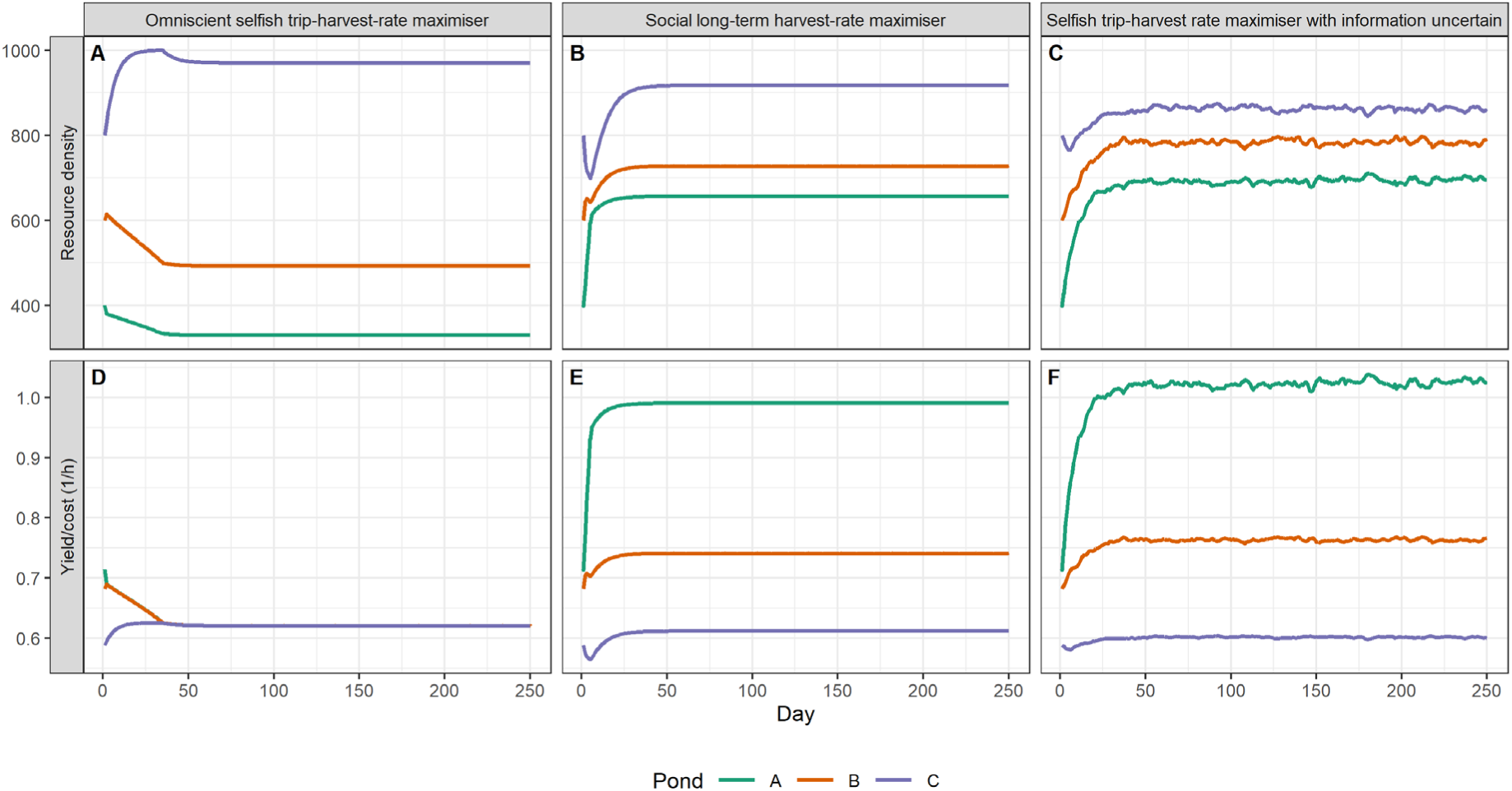
Changes in food density and yield per unit cost in each pond during the simulation run. For the omniscient trip-harvest-rate maximiser (A, D), the emerging yield per unit cost is similar for all ponds and relatively low (D). To increase the long-term yield per unit cost and minimize overexploitation, the harvester could take a social and more sustainable exploitation strategy based on the marginal gains (B, E). In that case, the yield per unit cost varies between ponds, but on average is substantially larger than the selfish trip-harvest-rate-maximiser (D). Alternatively, the harvester could maintain a selfish and short-term strategy, but when it is exposed to natural stochasticity and has limited cognitive abilities (e.g. only memorizing the last visit to each pond), the food density (C) and yield per unit cost (F) is similar to that of the social and long-term-harvest-rate maximiser (B and E).

**Figure 4.**
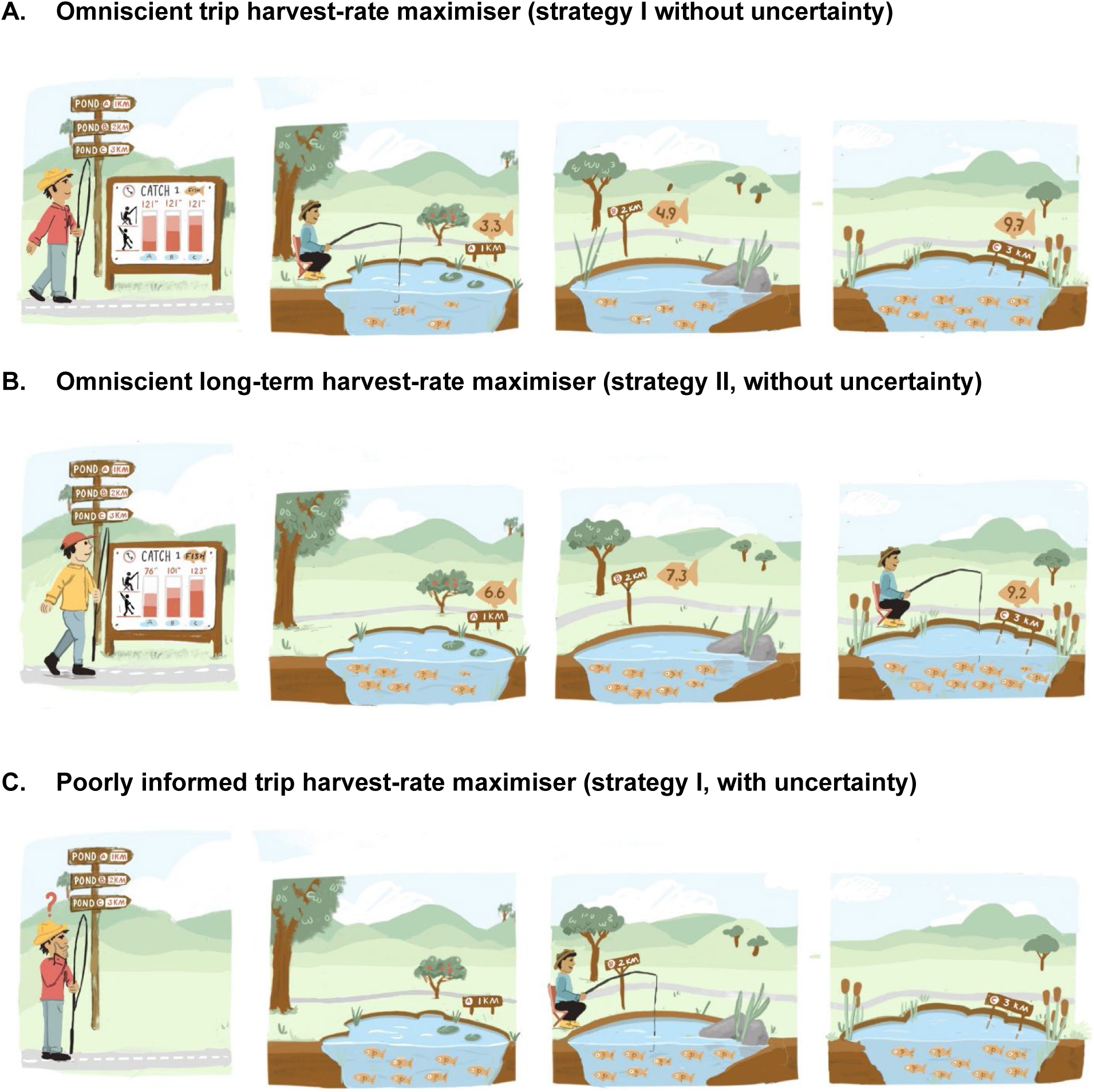
The emerging consequences of the two harvesting strategies (with and without uncertainty) on the equilibrium fish density and time required to harvest a single fish. When arriving at the initial state (Figure 1), the angler has the option of choosing between two harvesting strategies: either minimize the trip cost (Fig. 2A) or maximise the marginal gain (Fig. 2F) by taking fish productivity (Fig. 2C) into account. The illustrations above show the emerging equilibrium state for the omniscient trip harvest-rate maximiser (A), the omniscient sustainable-term harvest-rate maximiser (B) and the trip harvest-rate maximiser who exclusively relies on the memorised fishing duration experienced during the most recent visit to each pond (C). The information sign displays the total time (in minutes) required to catch a fish in each pond. This total time consists of the time spent walking (bottom bar in red) and fishing (top bar in pink). For scenario C, the harvester is unaware of the total time required to harvest a fish from each pond (73, 98 and 125 minutes, respectively). The number of fish in each pond (and the sign adjacent to the pond) depicts the fish abundance in each pond, with each fish illustration representing 100 individuals. For scenario C, the ponds contain 6.9, 7.8 and 8.6 fish, respectively, however the harvester is again unaware of these precise values. Harvesting conducted by omniscient trip harvest-rate maximisers results in a steep gradient in fish densities, with the lowest density occurring close to the harvester’s starting point, but the total harvest time will be equal across all ponds. In contrast, the sustainable harvest-rate maximiser aims to sustain higher fish density in each pond, leading to the shortest time required to harvest a fish. Under this strategy, the equilibrium cost of harvesting one fish will be significantly lower for the left pond than for the right pond, making the system more susceptible to defectors who aim to minimize trip costs. When the harvester attempts to maximise the trip harvest-rate by relying solely on the memorized catch rate from the most recent visit to each pond, the emerging fish density and time required to catch a fish in each pond will approximate those experienced by the sustainable harvest rate maximiser.

**Figure 5.**
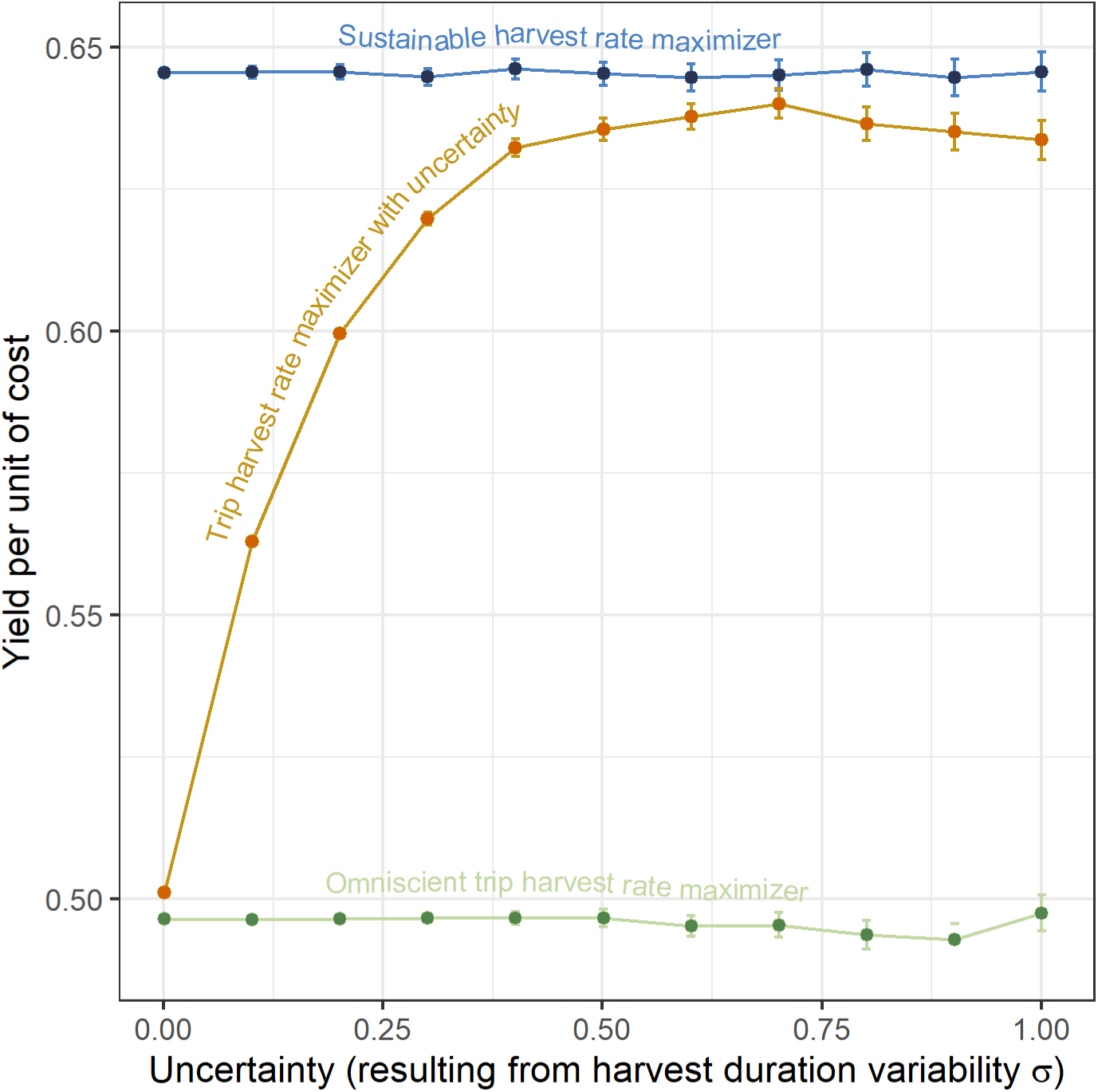
Variability in harvest duration (σ) and its impact on long-term yield per unit cost under various harvesting strategies. In the case of non-omniscient harvesters (orange line), this variability also leads to uncertainty in the perception and memory of resource density. The omniscient trip harvest rate maximisers and sustainable long-term harvest rate maximiser are also confronted with variability in harvest duration; however, they base their decisions where to forage on perfect information regarding resource density.

**Figure 6.**
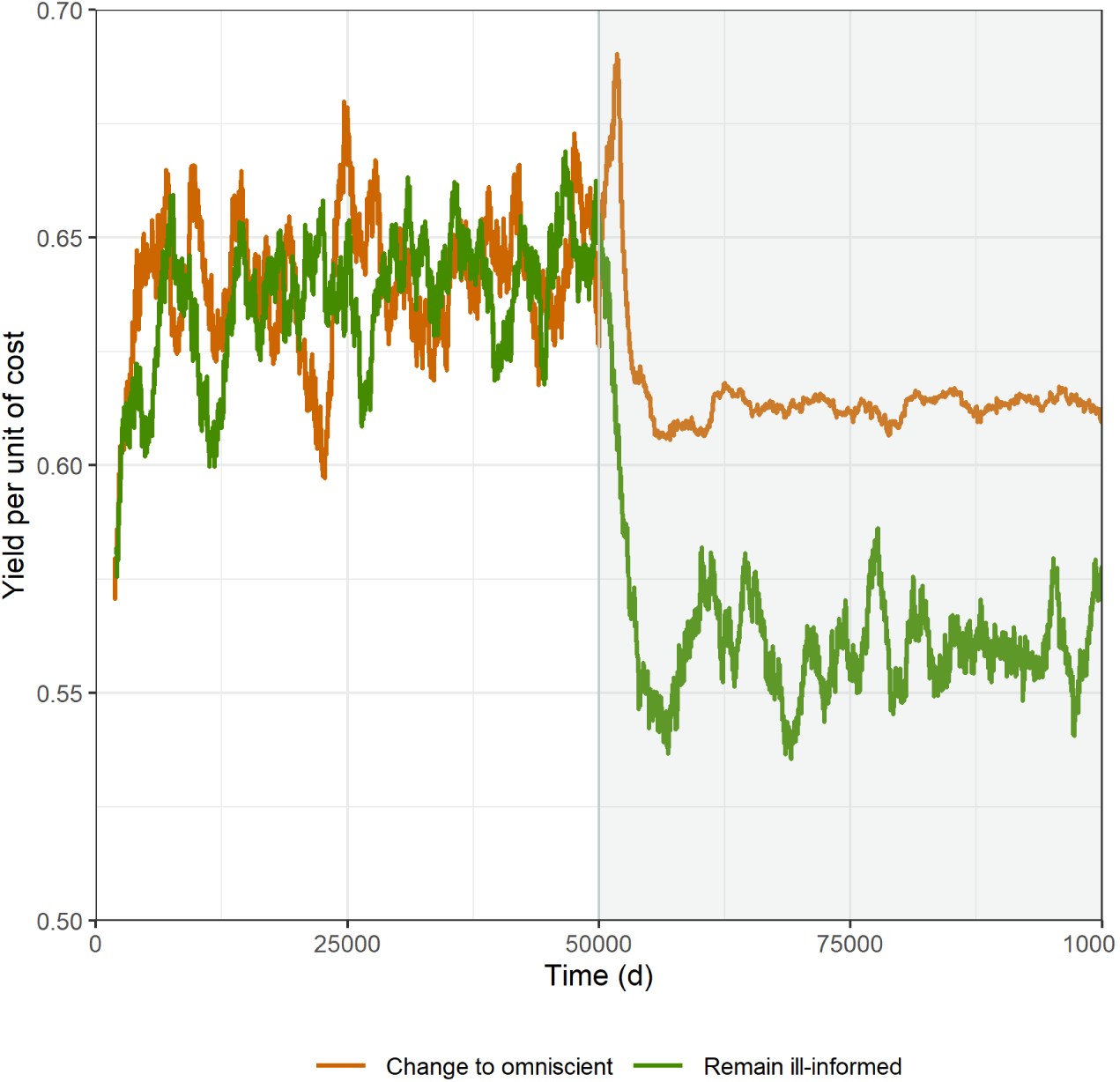
The effect of ‘invasion’ of omniscient individuals on the long-term yield per unit cost. There are two groups of individuals that are mixed and in competition, with each group comprising 25 individuals each. At the onset of the simulation, all individuals face uncertainty regarding the density of resources. At the midpoint of the simulation, one group transforms into omniscient harvesters (represented by the brown line), which leads to an overall decrease in long-term yield per unit cost. However, among the two groups, the omniscient foragers achieve a higher yield per unit cost.

If the angler chooses instead to maximise the long-term yield per unit cost (eq. 3), the equilibrium marginal costs will be equal for all ponds. To achieve higher yield, the harvester will first target location C (note the dip in Fig. 3c), and allow resources to renew elsewhere, particularly at pond A, which has the lowest initial resource density. Eventually, the resource density in all ponds will increase relative to their initial condition, and the resource density will be (well) above MSY for all ponds (Fig. 3C). The emerging trip duration will be close to 2.0h for the most distant pond (C) but compared to the short-term harvest-rate-maximiser, substantially lower at pond A and B, namely 1.26h and 1.69h, respectively (Fig. 3D). Hence, while the individual would regularly have to make more costly trips (to pond C), in the long-term they will be better off because the average cost per unit of yield will be lowest overall (1.5 h, relative to 2.0h for the trip-rate-maximiser).

### Harvesting with uncertainty

If the angler attempts to maximise the short-term yield per unit cost but is unable to make accurate judgements about the resource density in each pond due to stochasticity in catch duration, the long-term yield per unit cost can be substantially different. If the stochasticity is very small (σ = 0.001), the long-term yield per unit cost will be roughly equivalent to the omniscient harvester. The minor difference between the outcomes of the two strategies results from the harvester with uncertainty being unaware of the resource renewal occurring overnight, whereas the omniscient harvester is aware. When the stochasticity in catch duration increases, so does the long-term yield per unit cost, and at high stochasticity (σ^2^ = 1), the long-term yield per unit cost approaches that of an omniscient long-term harvest maximiser. For pond A, B, and C, the emerging resource densities under a high uncertainty scenario (σ = 1) are 691, 780 and 859, respectively (Fig. 3C). These values are closer to densities observed for the sustainable harvest rate maximiser (655, 727 and 917, respectively, see Fig. 3B), compared to the values observed for the omniscient trip harvest rate maximiser (329, 492 and 970, respectively, see Fig. 3A).

Figure 5 illustrates the simulation outcomes for a given set of parameters and demonstrates that a short-term maximiser faced with high uncertainty (σ = 1) can have a yield per unit cost close to that of a long-term maximiser, and substantially higher than that of an omniscient short-term maximiser. However, this outcome might be influenced by the chosen combination of parameter values. Appendix A, Fig. A3 illustrates the effect of different parameter values for the renewal rate (*r*), travel speed (*S*), and harvest rate parameter (*a*) on the performance of the different harvest strategies. If the rate of renewal *r* is low, the daily harvest exceeds renewal, resulting in the individual having to allocate an increasing amount of time to the harvest. This will ultimately lead to the exhaustion of the resource and eventual starvation of the consumer. In our simulation, this event occurred when the rate of resource renewal was set to 0.001. Depletion and collapse of the resource stock could also occur at a resource renewal rate of r = 0.0015, but only if both the harvest rate parameter (*α*) and travel speed (*S*) were low. When the resource renewal rates exceeded r = 0.002 (i.e., the value chosen for our main simulation), the resource renewal was sufficient to match the rate of resource harvest, at least within the parameter space explored. Under these conditions, the omniscient short-term maximiser experienced a lower yield per unit cost compared to the individual facing uncertainty, especially when the harvest rate parameter *α* was high, and the travel speed *S* was slow.

When examining the yield per unit cost of a short-term maximiser confronted with uncertainty, it approaches that of a long-term maximiser, especially at the lower resource renewal rates. The uncertainty scenario generally leads to higher long-term yields per unit cost compared to the omniscient scenario, except for situations involving high resource renewal, high travel speed, and low harvest rate (Appendix A, Fig. A3). In other words, when some overexploitation is likely to occur, the imperfectly informed short-term maximiser who is faced with uncertainty outperforms the omniscient one in the long term.

### Multiple harvesters and invasion

The simulation with multiple individuals allows for the incorporation of mixed strategies. Our focus was specifically on examining the change in yield per unit cost when some of the individuals facing uncertainty were substituted with omniscient individuals. At the onset of the simulation, all individuals maximised the short-term harvest rate while facing uncertainty. Halfway through the simulation, one group (50% of individuals) was replaced with omniscient individuals. After the ‘invasion’, all individuals (both the omniscient and uncertain ones) experienced a decrease in the yield per unit cost (Fig. 6). This decline in yield was most pronounced for individuals who were faced with uncertainty; thus, the omniscient harvesters outperformed the others in the mixed setting. However, the population as a whole achieved a much lower yield per unit cost.

## DISCUSSION

Competition by pre-emption for limited resources gives rise to the ‘tragedy of the commons’ [2,25]. This can lead to overexploitation, and in some cases, to the collapse of resources and their consumers [3,4]. If instead, all individuals take resource productivity into account and aim to maximise their long-term harvest, a sustainable and higher harvest situation can be achieved. However, this strategy necessitates perceptual, cognitive, and cooperation abilities that are unrealistic for nonhuman animals, and often challenging for humans [30]. Furthermore, even with modern technology, a perfect knowledge of resources’ dynamics is unrealistic, even for humans.

Here, we reveal that in nature there might exist a simpler solution to prevent overexploitation. When individuals prioritizing their own short-term interest but have imperfect knowledge of their environment, this can lead to a more sustainable harvest of resources, which is close to that expected for populations of perfectly informed individuals maximizing long-term harvest. This pattern was consistently observed across a wide range of parameters (Appendix A, Figure A3). So, while there might be a selective drive for individuals to make well-informed decisions as assumed by optimal foraging theory [31], consumers could be better off if they all make less well-informed decisions. This finding might explain why sub-optimal consumers and apparently sustainable exploitation are prevalent in nature, and why many experimental and field observations appear to contradict the predictions of optimal foraging theory [32–35].

So why does uncertainty in perceiving and memorizing resource density result in a higher long-term yield per unit cost? The reason is that misjudgements about the spatial distribution of resources lead to increased exploitation in remote and high-cost regions. Although making such misjudgements may lead to reduced cost-efficiency for a given trip, they are beneficial in the long run for two reasons. First, exploitation in remote areas can boost resource renewal there because these remote areas are often underexploited and well above MSY. Second, remote exploitation will also alleviate harvest pressure in nearby, low-cost harvesting areas with resource densities below MSY, which can lead to increased resource renewal also in these locations. Apparently, these uncertainty-induced misjudgements change the harvesting distribution such that long-term harvest rates become very similar to the ideal long-term strategy, which can typically only be achieved by fully cognitive and cooperative groups [21]. While environmental stochasticity is usually considered a risk factor for population viability [36], here we find theoretical evidence to suggest that populations living in stochastic environments may be forced to exploit resources more sustainably.

The absence of these counter-intuitive results in previous studies is likely due to the fact that the majority of research on movement and foraging strategies tends to focus on *individual-level* consequences of being confronted with imperfect information [37,38]. Indeed, also our results show that in a mixed setting, those individuals that do behave ‘optimally’ and maximise short-term harvest-rates, outcompete other individuals following a more sustainable strategy (Fig. 6). Only when most individuals deviate from short-term optimal behaviour, for example because uncertainty is causing them to do so, do individuals from that population attain higher long-term harvest rate. Historically, empirical support for individuals making such sub-optimal foraging movement decisions that contradict the optimal foraging theory has been limited, possibly due to a publication bias [39]. However, there is a growing body of literature on behaviours that appear maladaptive [40,41]. For example, an increasing number of empirical behavioural studies suggest the presence of sub-optimal decision-making, specifically studies related to gambling experiments. Pigeons have been observed exhibiting mal-adaptive behaviour by not selecting the option with the highest average food, but instead by choosing a risky option with a low probability of a high payoff [42]. Monkeys are also inclined to offer a higher price for information about a low probability, high-stakes gamble [43]. Humans who are prone to gambling exhibit similar tendencies, as they are willing to incur high costs to obtain non-instrumental information [44]. Additionally, they tend to place higher value on rewards when they have to exert more effort to obtain them, a phenomenon known as ‘justification of effort’ [45]. These findings suggest that humans and other non-human animals attribute an intrinsic value to information that does not align with optimal decision-making in uncertain situations but instead motivates them to engage in exploratory and information-seeking behaviour [46]. The work presented in this study shines new light on the potential evolutionary significance of such seemingly erroneous decisions.

### Future directions: Increasing complexity

The simulations in this study are intentionally simplistic: Our analysis considers a three-pond system with a single resource and a single type of central-place harvester, which has a fixed loading capacity, and is solely concerned with travel and harvesting costs. While this study specifically concentrates on central-place foragers, ultimately almost all animals have home ranges during certain periods of their life history [47]. And even more generally, as no organism on Earth has infinite movement capacity, travel costs are always involved. It is also worth noting that the potential benefits of information uncertainty may extend to harvesters who face spatially-varying costs other than or in addition to travel costs. These costs may include perceived predation risks (landscape of fear, e.g., [48]), unsuitable environmental conditions (e.g., [49]), or reduced prey catchability (e.g., [50]). Due to uncertainty in the perception of resource density, and potentially in associated costs or risks of exploitation, the harvesters may sometimes perceive a (slightly) suboptimal patch as being more valuable than it really is. By choosing to harvest in those patches, other patches may be left (temporarily) untouched, enabling resources to renew. Additionally, in our simulations, only one species is considered, which makes the system highly susceptible to overexploitation occurring in cyclic patterns [51]. In more natural systems, multi-species processes, such as top-down trophic control and competition among species, are among the pathways outlined by Vuorinen *et al.* [13] that counteract the evolutionary drive towards such overexploitation. If multiple species (e.g., other meso-predators) are competing for the same resources, or if other renewal functions such as the semi-chemostat resource-renewal functions [52,53] or sublinear renewal functions [54] occur, our observed positive effect of uncertainty on long-term harvest rate might no longer hold true. Finally, we considered a three-pond system, as opposed to a continuous heterogeneous landscape. Replicating the simulations for a continuous terrain in which the forager distribution extends in all directions would result in a quadratic increase in harvesting surface area with harvesting range, as opposed to a linear one. Consequently, in a two-dimensional landscape, we expect that individuals pursuing a short-term strategy may experience even greater benefits from uncertainty.

### From theory to societal implications

An important question that remains unresolved is why the occurrence of overexploitation is so prevalent among humans [55], while it is uncommon among many other animal species [13,55]. The primary cause of overexploitation by humans is attributed to their exceptional efficiency through the use of tools and advanced technology, as well as their cooperative hunting abilities and extensive mobility unparalleled among other predators [56]. In our study, we accordingly showed that overexploitation is more likely to occur when both harvest efficiency is high and uncertainty is low (Appendix A Figure A3). Typically, in other natural systems with such a superior consumer, excessive exploitation would lead to the depletion of the resource, followed by a collapse of the consumer population. Humans might have circumvented these negative feedbacks by continuously shifting to different, typically smaller, prey species or other natural resources [57]. Moreover, historically, uncertainty might have compensated for some of the increase in the efficiency and prevented overexploitation, but now, our technology, survey abilities and cartography has improved so immensely that this may no longer be the case.

Despite humans’ efficiency and intellect, sustainable exploitation (e.g. as proposed by Ling and Milner-Gulland [21]) is rarely observed [16]. What practical solutions can be employed to break free from this apparent inevitability? Achieving a more sustainable exploitation by collectively reducing our harvesting efficiency and collective memory seems to be practically infeasible. Thus, two avenues appear to remain. Social groups may adopt a long-term exploitation strategy predicated on the marginal gains theory as described here and in [21]. However, this would necessitate accurate information regarding the present and future productivity and distribution of resources, as well as an ability to restrain defectors. Although feasible in certain contexts, such as aquaculture or agriculture, it is improbable that the necessary information will ever be available for common pool resources extracted from natural systems. The second course of action is to recognize that it may be impossible to adopt an MSY strategy, and instead, to harvest less. In our simulation, consumers were assumed to harvest a fixed quantity of resources. However, spatial and temporal uncertainty in harvest rates may also influence how much groups of individuals harvest [58]. Indeed, a study examining the decision-making processes of real fishing communities showed that groups that are uncertain about thresholds are more likely to maintain higher stock levels, which could potentially prevent regime shifts [59]. Social security can also have an important influence on how much resources are extracted, because intergroup sharing, cooperation and communication can reduce the likelihood of resource collapse [60,61]. Clearly, experimental research on collective action related to resource extraction in the face of environmental and social uncertainty is highly relevant in light of the biodiversity and climate crises [62]. In the future, the connection of empirical studies with mechanistic modelling studies, such as the present one, should be at the forefront of research to ascertain the practical applicability of these mechanisms in real-life, human scenarios.

## Supporting information

Simulation function R-code

R-code for simulations

R-code for figures

## Acknowledgements

We thank Malou Zuidema for making the drawings shown in Fig. 1 and 4, and Jan van Gils and Eelke Folmer for reading a final version of the manuscript.

## Author contributions statements

GA, LR-L, JF, FvL and JM conceived the ideas and designed the methodology; GA led the analysis and simulation with support from LR-L and JF; GA and LR-L led the writing. All authors contributed critically to the text and gave final approval for publication.

## Data, scripts, code, and supplementary information availability

Scripts are appended as supplemental files to the BioRxiv repository. Files included are Simulation_function_BioRxiv_2025_01_15.R containing the main function used for the simulations, Simulations_3pond_BioRxiv_2025_01_15.R containing R-code to execute the simulations, and Result_plotting_BioRxiv_2025_01_15.R containing R-code for making the figures.

## Appendix A: Supplementary tables and figures

**Figure A2.**
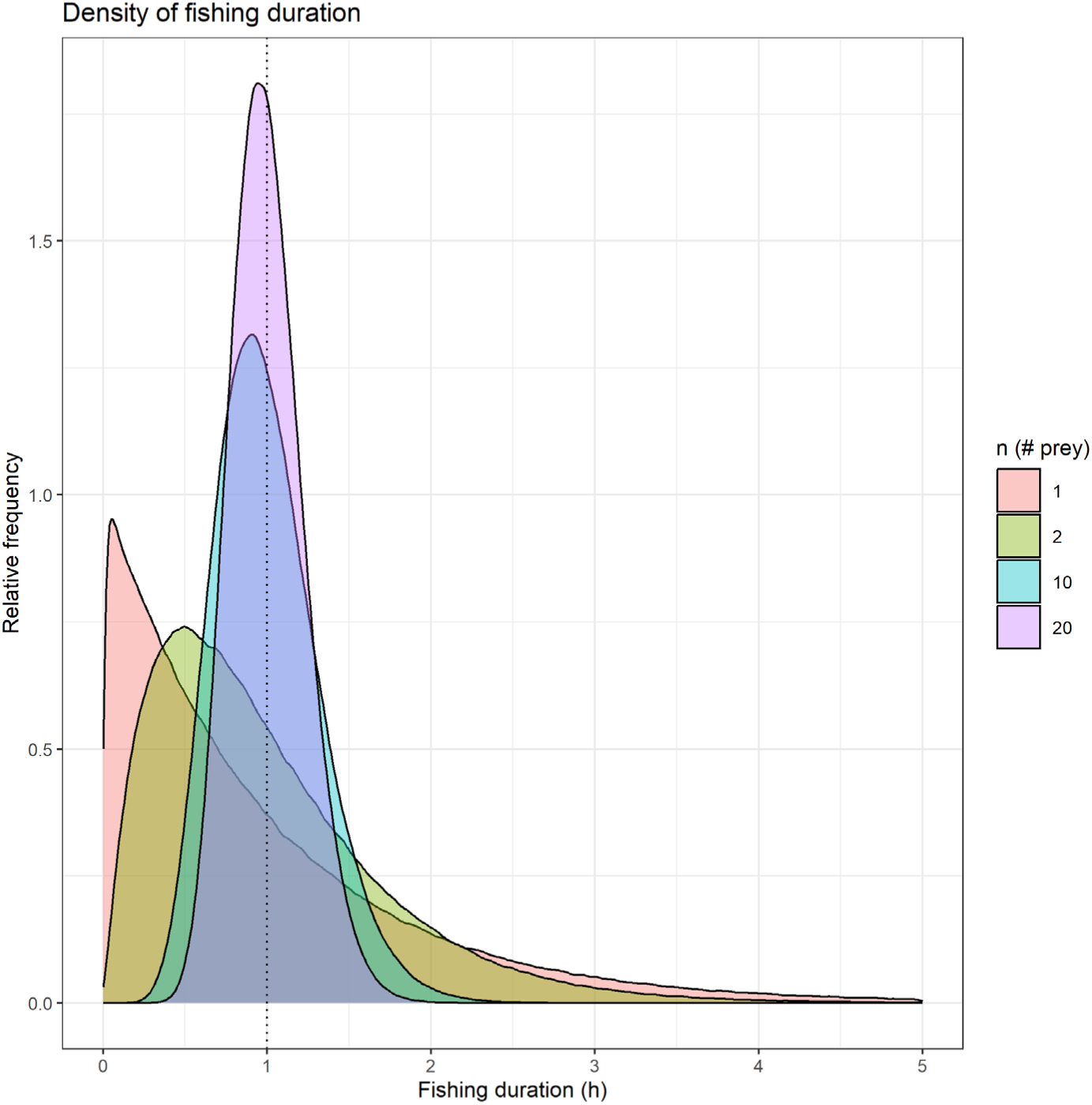
The effect of the number of fish caught (*η*) on the uncertainty in fishing duration assuming a gamma distribution. Increasing *η* (and proportionally increasing the fish density), the average harvest duration remains the same, but the variability in harvest duration decreases.

**Figure A3.**
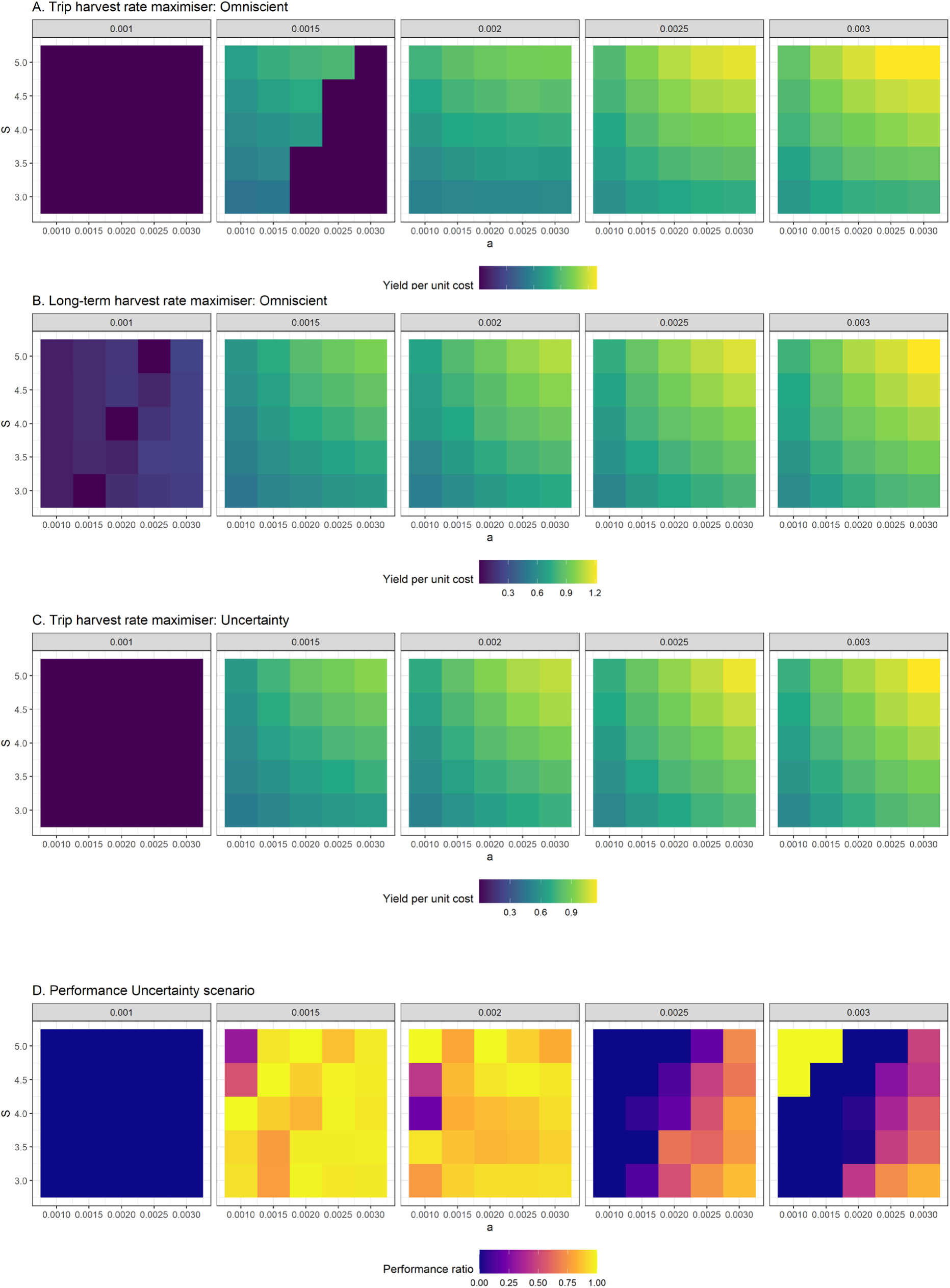
The effect of parameter values on yield per unit of cost. The parameter values include the travel speed (S, on y-axis), the harvest rate parameter (a, on x-axis) and resource renewal rate (r, panels). The performance ratio in figure D is defined as 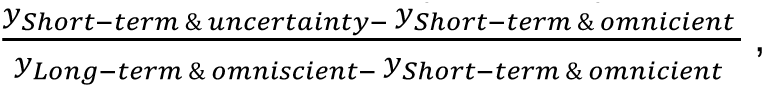 where y represents the yield per unit of cost. If this performance ratio is close to 1, the uncertainty scenarios lead to long-term intake rates which are almost equivalent to the maximum achievable long-term harvest strategy.

---

**Table A1.** Model parameters and baseline values used for generating figure 2.

| Harvest parameters | Symbols | Baseline values |
| --- | --- | --- |
| Cost of travelling one unit of distance (i.e. the inverse of travel speed $S$) | $\tau$ | 0.25 |
| Travel speed (in km/h) | $S$ | 4 |
| Cost of capturing one resource unit at resource density $n = 1\text{m}^{-2}$ (i.e. the inverse of the catch rate) | $\gamma$ | $1/0.002$ |
| Catch rate or the area harvested per unit of time (in $\text{m}^2/\text{h}$) | $\alpha$ | 0.002 |
| Resource parameters |  |  |
| Intrinsic resource population growth rate | $r$ | 0.002 |
| Local resource carrying capacity density | $k$ | 1000 |
| Spatially varying variables |  |  |
| Resource density at location $x$ | $n(x)$ | |
| Distance between the starting point and location $x$ | $d(x)$ | |
| Rate of resource productivity (and offtake at equilibrium) per unit of area | $q(n)$ | |

